# Convex representation of metabolic networks with Michaelis-Menten kinetics

**DOI:** 10.1101/2023.01.17.524421

**Authors:** Josh A. Taylor, Alain Rapaport, Denis Dochain

## Abstract

Polyhedral models of metabolic networks are computationally tractable and can predict some cellular functions. A longstanding challenge is incorporating metabolites without losing tractability. In this paper, we do so using a new second-order cone representation of the Michaelis-Menten kinetics. The resulting model consists of linear stoichiometric constraints alongside second-order cone constraints that couple the reaction fluxes to metabolite concentrations.

We formulate several new problems around this model: conic flux balance analysis, which augments flux balance analysis with metabolite concentrations; dynamic conic flux balance analysis; and finding minimal cut sets of networks with both reactions and metabolites. Solving these problems yields information about both fluxes and metabolite concentrations. They are second-order cone or mixed-integer second-order cone programs, which, while not as tractable as their linear counterparts, can nonetheless be solved at practical scales using existing software.

## 1. Introduction

The structure of a metabolic network contains useful information about its cellular functions. Two techniques for analyzing this structure are

- flux balance analysis (FBA), in which optimization is used to predict reaction fluxes [1], and
- minimal cut set (MCS) analysis, which attempts to find critical subsets of reactions that, when removed, disable certain functions [2].

Standard formulations of FBA and MCS analysis are based on a linear approximation in which only the reaction fluxes are variables. The benefit of this simplification is that FBA is a linear program (LP), which can be reliably solved at large scales, and powerful analytical tools like Farkas’ Lemma are available for finding the MCSs of polyhedral systems. Quoting [1], “FBA has limitations, however. Because it does not use kinetic parameters, it cannot predict metabolite concentrations.” For the same reason, MCS analysis cannot explicitly identify critical metabolites.

In this paper, we augment FBA and MCS analysis with Michaelis-Menten kinetics and metabolite concentrations. Using the results of [3], we represent the Michaelis-Menten kinetics as a second-order cone (SOC) constraint. This leads to several original problem formulations.

- *Conic FBA (CFBA)*. CFBA predicts both reaction fluxes and metabolite concentrations in steady state. It is a single SOC program (SOCP), which, while not as tractable as LP, can be solved at practical scales [4]. We also formulate dynamic CFBA, in which the SOC representations of the reaction kinetics are used in dynamic FBA [5].
- *Conic MCS (CMCS)*. A CMCS is a cut set through a network of paths from reactions to metabolites, as specified by the stoichiometric matrix, and from metabolites to reactions, as specified by the Michaelis-Menten kinetics. To solve for CMCSs, we follow the strategy of [6], which uses Farkas’ Lemma [4] and results from [7, 8] on irreducible infeasible subsystems (IIS) to identify MCSs. We generalize this to CMCSs using the recent results of [9] on the IISs of semidefinite systems. We also formulate a linear approximation that, due to the discrete nature of cut sets, produces similar results.

We remark that CFBA and CMCS analysis might not produce better predictions of reaction fluxes than existing methods. The main benefit of CFBA and CMCS analysis is that they incorporate metabolite concentrations and reaction kinetics while retaining much of the tractability of standard linear formulations.

We now describe how our contributions relate to the existing literature. Using LP to analyze metabolic networks was first suggested in [10]. Since then FBA has gained wide acceptance [11, 1], and is available in open source implementations [12, 13]. Dynamic FBA (DFBA) is an extension that incorporates reaction kinetics, which are typically nonlinear, and metabolite concentrations, potentially as well as other transient information such as reprogramming and light intensity. Reference [5] first formulated the two main types of DFBA. The ‘dynamic optimization approach’ is a large nonlinear program, in which the reaction kinetics constrain the fluxes through time. In the ‘static optimization approach’, one solves a sequence of LPs and integrates the solution between time periods. There have been several refinements such as putting the optimization directly into the ODE simulation [14], and using lexicographic optimization to improve robustness when the LP has multiple solutions [15].

CFBA is similar to DFBA in that it also captures metabolite concentrations and reaction kinetics, but only when modeled as Michaelis-Menten. They differ in that CFBA is in steady state, and hence does not capture transients. If one approximates the biomass concentration as a constant, or approximates Michaelis-Menten with the Contois function [16], one can formulate the dynamic optimization approach of [5] as an SOCP. We refer to this as Dynamic CFBA. Dynamic CFBA can capture transients and accommodate time-varying parameters. CFBA and dynamic CFBA are not necessarily more accurate than dynamic FBA, but are more tractable in that they are single SOCPs.

CFBA is also related to resource balance analysis (RBA) [17 18], a more general problem that can predict fluxes, metabo lites, macromolecular cellular processes, and proteins. RBA differs from CFBA in that it does not contain the Michaelis-Menten function or any other nonlinearities, and as a result is an LP. In principle, our SOC representation of the Michaelis-Menten function could be incorporated into RBA, leading to an SOCP.

Another relevant literature stream focuses on the analysis of pathways through metabolic networks (see, e.g., [19, 20]). In the linear case, the nonnegativity and stoichiometric constraints form a polyhedral cone, the extreme rays of which correspond to the elementary flux modes of the metabolic network [21]. It is not clear that this perspective extends to our setup because SOC constraints are non-polyhedral, and the SOC representation of the Michaelis-Menten kinetics is not in fact a cone. The Minkowski-Weyl Theorem states that a cone is finitely generated if and only if it is polyhedral [22]. Therefore, such a system could have an infinite number of elementary modes if it has any at all. If elementary modes do exist, at present we are not aware of any reliable techniques for obtaining them.

For these reasons, we focus on the adjacent problem of identifying MCSs. Reference [6] showed that the MCSs of a metabolic network are the elementary modes of a dual network specified by Farkas’ Lemma [4]. We make use of the results of [9] to generalize this strategy to networks with metabolite concentrations coupled to the reaction fluxes through Michaelis-Menten kinetics. By then using a linear approximation of the Michaelis-Menten kinetics, we recover the use of tools for polyhedral systems, which we find produce more reliable results.

The paper is organized as follows. Section 2 reviews metabolic network modeling and the SOC representation of the Michaelis-Menten kinetics. Section 3 presents FBA, CFBA, and dynamic CFBA. Section 4 presents the MCS analysis of [6] and our extensions to systems with Michaelis-Menten kinetics. In Section 5, we apply CFBA, dynamic CFBA, and CMCS analysis to a model of *Escherichia coli*.

## 2. Background

### 2.1. Metabolic networks

The reactions are indexed by the set 𝒩, where *n* = |𝒩|. Let ℛ ⊆ 𝒩 and ℐ ⊆ 𝒩 be the sets of reversible and irreversible reactions, where ℛ ∪ ℐ = 𝒩 and ℛ ∩ ℐ = ∅. There are a total of *m*′ metabolites.

Let *z* ∈ ℝ^*m*′^ be a vector of metabolite concentrations and *v* ∈ ℝ^*n*^ a vector of fluxes due to the reactions. *S* ∈ ℝ^*m*′×*n*^ is the stoichiometric matrix. The dynamics of the metabolites are given by

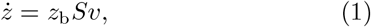

where *z*_b_ is the element of *z* corresponding to biomass. We note that while the biomass is not a metabolite, its evolution can be written as *z*_b_ times a linear combination of the reaction fluxes, and hence can be represented as a row of (1).

In quasi-steady state, *ż* = 0. We then have *z*_b_*Sv* = **0**, which, because *z*_b_ is a nonzero scalar, implies *Sv* = **0**. The fluxes are subject to the bounds 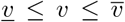. Typically, *<u>v</u>*_*i*_ = 0 for *i* ∈ ℐ and −∞ otherwise. When *<u>v</u>* = **0** and 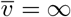, the system is a polyhedral cone [21].

Quasi-steady state models generally do not explicitly model the metabolites. Here we will include a subset of them, ℳ, where *m* = |ℳ|. We will henceforth let *z* ∈ R^*m*^.

Some of the fluxes are also bounded by a nonlinear function, usually the Michaelis-Menten kinetics [23]. We denote this set by 𝒬 ⊆ 𝒩, where *q* = |𝒬|. We expect that *q* ≥ *m*, because otherwise there are metabolites that appear in no reaction, and thus have no coupling to the rest of the model. To simplify notation, we order 𝒩 so that its first *q* elements are 𝒬. Let *V*^max^ ∈ ℝ^*q*^ and *K*^m^ ∈ ℝ^*q*^ be vectors of maximum reaction rates and Michaelis constants. We can write this bound for each reaction *i* ∈ 𝒬 as

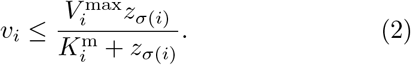

Here *σ*(*i*) identifies the index of the metabolite concentration appearing in reaction *i*. In general *σ* is not invertible because the same metabolite can appear in multiple reactions. Note that if *i* is a reversible reaction, (2) may be applied in the reverse direction by putting a minus sign in front of *v*_*i*_.

When a reaction has multiple reactants, its flux may be limited by more complicated functions such as the product of several Michaelis-Menten kinetics (cf. Table IV in [24]). We can straightforwardly generalize our notation to this case. Suppose that reaction *i* has *p*_*i*_ reactants. For *k* = 1, …, *p*_*i*_, let *σ*_*k*_(*i*) be the index of the *k*^th^ metabolite in the reaction. Then we can write the upper bound on reaction *i* ∈ 𝒬 as

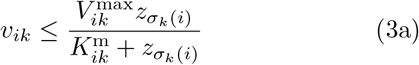

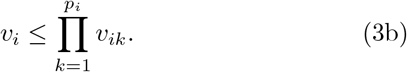

The intermediary variables, *v*_*ik*_, *k* = 1, …, *p*_*i*_, will be convenient for representing (3) in SOC form.

### 2.2. Convex representation of the Michaelis-Menten kinetics

As shown in [3], we can represent the inequality (2) as the SOC constraint

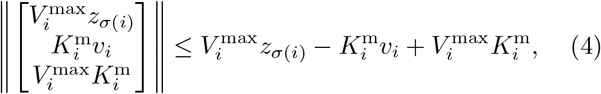

for *i* ∈ 𝒬. One can confirm equivalence by squaring both sides and simplifying. SOCPs consisting of linear and SOC constraints can be solved at large scales [4]. We provide a brief overview of SOCP in Appendix A.

Suppose now there are multiple reactants. As with (2), we can write (3a) as an SOC constraint in the form of (4). Unfortunately, (3b) is nonconvex. We can instead approximate it with

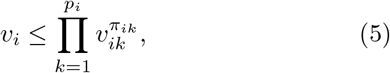

which has an SOC representation if 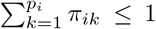 [25]. When *π*_*ik*_ = 1*/p*_*i*_ for *k* = 1, …, *p*_*i*_, it is the geometric mean. (5) is ad hoc in that it cannot in general be derived from first principles. However, we believe it could be a useful approximation because there is considerable room to tune the fit via the exponents and Michaelis-Menten parameters; and, similar ad hoc expressions have been used in the past when Michaelis-Menten alone was not sufficiently descriptive (cf. Table IV in [24]).

In the simple case where *p*_*i*_ = 2 and *π*_*i*1_ = *π*_*i*2_ = 1*/*2, (5) is a hyperbolic constraint with SOC representation

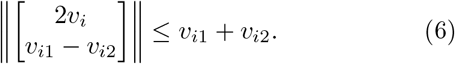

Sometimes LP is more practical to work with than SOCP, e.g., when there are numerous discrete constraints or commercial solvers are too expensive. Fortunately an SOCP can always be approximated to arbitrary accuracy with an LP, albeit a potentially large one. Reference [26] provides a constructive procedure for approximating a generic SOC constraint with a family of linear constraints. Alternatively, the right hand side of (2) can be straightforwardly approximated by a family of line segments.

## 3. Flux balance analysis

In this section, we first review the standard formulation of FBA, and then formulate CFBA and Dynamic CFBA. We also use the duals to derive several sensitivities.

### 3.1. Linear FBA

The below LP is a basic FBA routine.

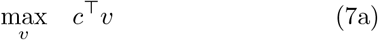

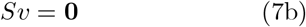

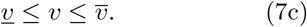

Here *c* ∈ ℝ^*n*^ selects and/or weights the fluxes for maximization. Let ℱ denote the optimal objective value.

Let *λ* ∈ ℝ^*m*^ be the vector of dual multipliers of (7b), and let 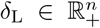 and 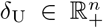 be the vectors of dual multipliers associated with the upper and lower bounds in (7c). The dual of (7) is

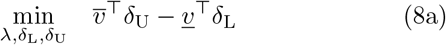

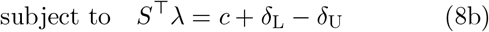

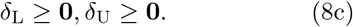

The dual variables, or shadow prices, can be interpreted as the sensitivities of the objective in (7) to changes in the constraints. More precisely, for each *i* ∈ 𝒩,

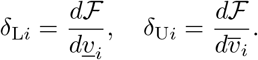

*λ* is similarly interpretable as the sensitivity of the objective to perturbations to (7b). This, for example, can be used to determine which reactions are most influential [27, 28].

We can also obtain insight by observing that the objectives of (7) and (8) must be equal due to strong duality:

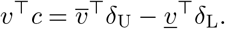

This breaks down the optimal objective into contributions from each constraint. In the common case where we maximize a single reaction, *v*_*s*_, and *v* = **0**, this simplifies to 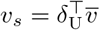, a convex combination of the upper flux bounds.

### 3.2. Conic FBA

We now generalize the FBA (7) to include metabolite concentrations, which are coupled to the reactions by the Michaelis-Menten kinetics.

Let *z* ∈ ℝ^*m*^ and 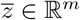 be vectors of upper and lower bounds on the metabolite concentrations. Let *d* ∈ ℝ^*m*^ be a vector weighting the metabolites in the objective.

Consider the below optimization.

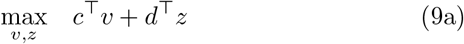

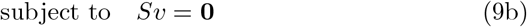

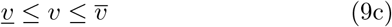

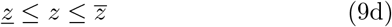

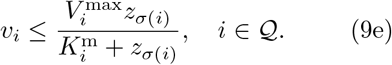

We make the following observations.

- This is an SOCP if we write (9e) in the form of (4).
- (9e) is only meaningful if the corresponding metabolite concentrations are minimized in the objective, i.e., *d <* 0. Otherwise, it is trivially optimal to fix the metabolites at their maximum concentrations, 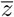, in which case (9e) can be represented as an upper bound on *v* in (9c).
- (9) allows for both internal and external metabolites [29]. However, if the external metabolites are buffered, which is to say fixed in our model, then the corresponding Michaelis-Menten function in (9e) is also fixed, and is then more efficiently represented as part of the vector 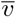.

We now derive the dual. For matrices *A* and *B* of the same size, let *A* ∘ *B* denote the element-wise product. We let *A*_*i*:_ and *A*_:*j*_ denote the *i*^th^ row and *j*^th^ column of *A*.

Let *λ* ∈ R^*m*^ be the vector of dual multipliers of (9b), 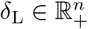 and 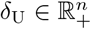 of (9c), and 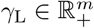 and 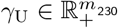 of (9d).

Let Φ ∈ ℝ^3×*q*^ and 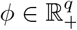. For a given reaction *i* ∈ 𝒬, the constraint (9e) (in SOC form (4)) has dual multipliers 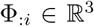 and *ϕ*_*i*_ ∈ ℝ_+_, which satisfy the SOC constraint ∥Φ_:*i*_∥ ≤ *ϕ*_*i*_.

Let *M* ∈ ℝ^*q*×*m*^ be such that *M*_*ij*_ = 1 if *σ*(*i*) = *j* and zero otherwise. The interpretation of *M* is complementary to that of *S. S* encodes a network representing how the reactions influence the evolution of the metabolite concentrations. Similarly, *M* encodes a network representing which metabolites appear in each reaction.

The dual of (9), which is also an SOCP, is below.

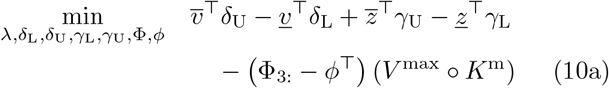

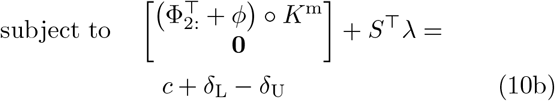

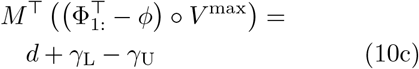

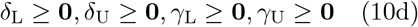

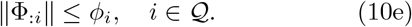

Strong duality holds if a constraint qualification is satisfied, e.g., Slater’s condition [4]. In this case *λ, δ*_L_, *λ, δ*_U_, *γ*_L_, and *γ*_U_ all have the usual LP sensitivity interpretations and complementary slackness with their respective constraints.

We can similarly interpret Φ and *ϕ*; see, e.g., Section 5.9.3 of [4]. *ϕ*_*i*_ is the sensitivity of the optimal objective value, ℱ, to perturbations to the right hand side of (4). Φ_:*i*_ is a vector of sensitivities of ℱ to perturbations to each element in the left hand side. We can use the chain rule to derive sensitivities for the parameters of the Michaelis-Menten function. For *i* ∈ 𝒬 we have

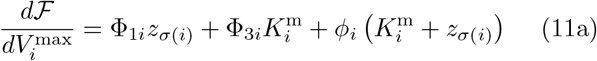

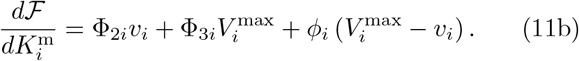

As in the previous section, the objectives of (9) and (14) match if strong duality holds. If we assume that *<u>v</u>* = **0**, *<u>z</u>* = **0**, and 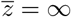, the equality simplifies to

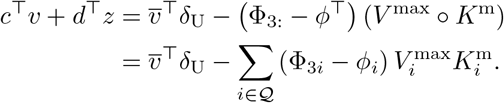

#### 3.2.1. Alternative objectives

A shortcoming of (9) is that the value of the optimal objective—flux rates minus metabolite concentrations— does not have a clear physical meaning. We could instead maximize *d*^⊤^*z* alone subject to a constraint on *v*, e.g., *v*_*s*_ ≥ 1, where *s* is the index of a reaction of interest. This is interpretable as the minimum metabolite concentration necessary to carry out a certain function. Such a constraint could be incorporated into (9c).

An objective consisting only of metabolite concentrations might be more physically interpretable. For instance, minimizing metabolite concentrations was used as a model of cellular function in [30]. A second advantage is that the dual sensitivities in (11) also have physical meanings. For these reasons, we maximize −**1**^⊤^*z* in the example in Section 5.1.

We could also maximize *c*^⊤^*v* alone and add constraints on *z*. This would be nontrivial, e.g., with coupled polyhedral constraints. However, box constraints like (9d) would simply lead to *z* binding at its upper bounds.

Note that while the dual is slightly different for these alternative formulations, the expressions for the sensitivities to *V*^max^ and *K*^m^ in (11) are unchanged.

### 3.3. Dynamic FBA

Dynamic FBA extends conventional FBA in two ways: the incorporation of metabolite concentrations, and changes over time not captured in a steady state model. Metabolite concentrations are present in CFBA, but not transients. We thus formulate dynamic CFBA for the purpose of capturing transient phenomena.

There are two main approaches to dynamic FBA, as described in [5].

- In the dynamic optimization approach, the full trajectory is optimized. The concentrations across time periods are coupled by a finite difference approximation of the derivative; note that we could use a more accurate approximation, e.g., a higher order Runge-Kutta scheme [31]. The optimization is nonlinear due to the Michaelis-Menten kinetics and bilinearities between the fluxes and biomass variables.
- In the static optimization approach, a conventional FBA is solved in each time period. The resulting flux vector is used to propagate the concentrations to the next time period.

The first is harder to solve because it is larger and nonlinear. Here, we optimize the full trajectory as in the dynamic optimization approach. We render the problem more computationally tractable by representing the Michaelis-Menten kinetics in SOC form, (4).

A difficulty not present in steady state FBA is the biomass, *z*_b_, which we recall is an element of the vector *z*. Because we are not setting the derivatives in (1) to zero, we cannot divide it out, and the product of the Michaelis-Menten function and *z*_b_ does not have an SOC representation. We can remedy this in two ways.

1. If the biomass evolution is predictable, we can approximate it as a fixed parameter in each time period, 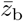(*t*). Then the kinetics can be represented in the form of (4).
2. Instead of Michaelis-Menten, we can model the reaction limits with the Contois function [16]:

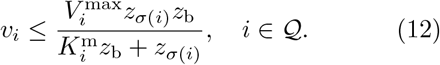

The Contois function is commonly used to model biochemical processes [32], and differs from Michaelis-Menten only in that the denominator depends on *z*_b_. Given its similarity to the Michaelis-Menten kinetics, (12) could also be an acceptable approximation. The advantage of the Contois function is that, unlike the product of *z*_b_ and the Michaelis-Menten function, the resulting kinetics has an SOC representation. As shown in [3], we can represent the inequality (12) as the SOC constraint

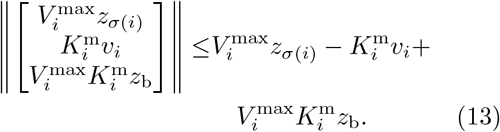

Which approximation is more appropriate depends on the context. For example, if the biomass does not vary much, holding it constant in Michaelis-Menten is a clear choice; one can test this assumption by subsequently simulating (1). If the evolution of the biomass does depend strongly on the the other metabolites, then the Contois function might be a better choice. Alternatively, we could fit a generic SOC constraint to reaction data, which would retain compatibility with SOCP and potentially be more accurate than either of the above approximations [33].

There are time periods *t* = 0, …, *τ*, each of length Δ. Let *v*(*t*) and *z*(*t*) denote the fluxes and metabolite concentrations in period *t*. We thus obtain the below SOCP for dynamic CFBA.

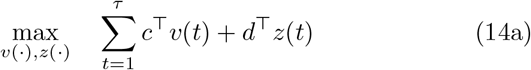

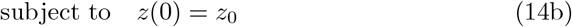

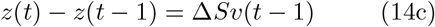

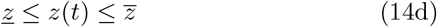

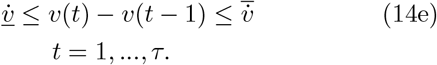

If we approximate the biomass as a fixed parameter, we also have for *t* = 1, …, *τ* :

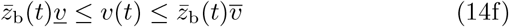

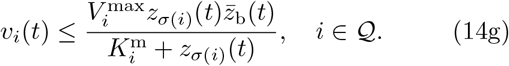

If we model the reaction limits with the Contois function, (12), then instead of (14f) and (14g), we have for *t* = 1, …, *τ* :

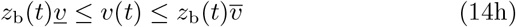

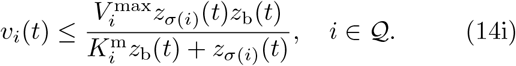

(14b) is the initial condition. (14c) is the Euler approximation of (1), and couples the variables across time periods. (14d) and (14f) (or (14h)) are bounds on the reactions and metabolites. (14e) limits the rate of change of the reactions. In (14g) (or (14i)), *v*_*i*_(*t*) is less than either the Michaelis-Menten kinetics with fixed biomass or the Contois function, depending on which of the above approximations is used.

Note that in dynamic FBA, the evolution of the metabolites generally depends on more than just the stoichiometry, as in (14c). The same should also be the case for dynamic CFBA. For example, there could be inflows and outflows with endogenous or exogenous metabolite concentrations, or biomass death [15].

## 4. Minimal cut sets

In (11) in Section 3.2, we used dual variables to compute sensitivities to kinetic parameters. This was one of several potential applications of the CFBA dual system. We now explore another in which we use Farkas’ Lemma to identify CMCSs.

An MCS is the smallest set of reactions which, if constrained to be zero, disables some function of interest. In this regard, the reactions in an MCS are the lynchpins of the system. In [6], it was shown that the MCSs of a metabolic network correspond to the elementary modes of a dual network, which is specified by Farkas’ Lemma [22]. This was an application of a result from [7, 8], which states that there is a one-to-one mapping between the IISs (irreducible infeasible subsystems) of a linear system and the vertices of its dual polyhedron. One can therefore identify the MCSs of a metabolic network by solving for the vertices of the dual polyhedron, which can be done via mixed-integer LP (MILP).

In [9], a weaker version of the result of [7, 8] was extended to semidefinite systems, which generalize SOC systems. We make use of this in Section 4.1 to extend the results of [6] to networks with metabolites linked by Michaelis-Menten kinetics. Whereas an MC contains only reactions, a CMCS contains the smallest set of reactions and/or metabolites that the system cannot function without. We then formulate a linear approximation that produces similar results in Section 4.2.

### 4.1. Conic minimal cut sets

We now describe the setup, starting with the linear part as given in [6]. Let 𝒯_*v*_ ⊆ 𝒩 denote the polyhedral set of target reactions, parametrized by the matrix *T* and vector *v*^*^.The target set encodes some function of interest, which the removal of a cut set disables. The constraint

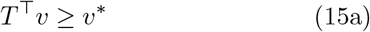

forces the reactions in 𝒯_*v*_ to be active. The below two constraints encode the steady state operation of the metabolic network:

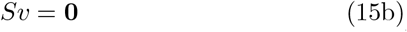

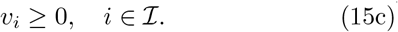

We say that *C*_*v*_ is a cut set for 𝒯_*v*_ if *v*_*i*_ = 0 for *i* ∈ *C*_*v*_ implies that *v*_*i*_ = 0 for *i* ∈ 𝒯_*v*_ under (15b) and (15c). It is an MCS if it contains no smaller cut sets, i.e., cut sets with fewer elements, for 𝒯_*v*_.

To make (15a)-(15c) infeasible, following [34], we add the constraints

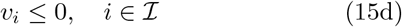

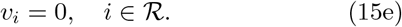

Note that together, (15c)-(15e) imply *v* = **0**.

Lemma 1 in [6] states that each MCS corresponds to an IIS of (15a)-(15e). Note that an MCS could correspond to multiple IISs, but no two MCSs correspond to the same IIS.

We now incorporate metabolite concentrations and reaction kinetics. We assume there is also a target set of metabolites, 𝒯_*z*_, each of which is constrained to be in some concentration range by

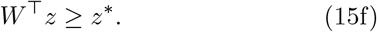

The below two constraints prohibit negative concentrations and relate the concentrations to the reactions:

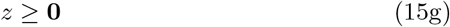

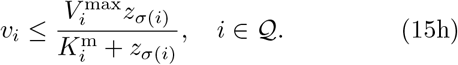

Similar to (15d) and (15e), we make this portion of the system infeasible by adding the constraint

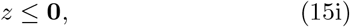

which, together with (15g), implies *z* = **0**.

We note that infeasibility is not always precise enough when moving from linear to nonlinear systems because in the latter case, the set of solutions might not be closed. As in [9], we say that a semidefinite system is weakly feasible if any positive perturbation to its eigenvalues makes it feasible, and weakly infeasible if it is not weakly feasible. Feasibility implies weak feasibility, and weak infeasibility implies infeasibility. (15) is weakly infeasible, and hence also infeasible, because there are positive perturbations that do not make it feasible. We henceforth take the definition of an IIS to refer to weak infeasibility.

We now extend the definition of an MCS.

#### Definition 1

**𝒞** = {𝒞_*v*_, 𝒞_*z*_} *is a cut set for* {𝒯_*v*_, 𝒯_*z*_} *under (15b), (15c), (15g), and (15h) if the additional constraints v*_*i*_ = 0 *for i* ∈ 𝒞_*v*_ *and z*_*i*_ = 0 *for i* ∈ 𝒞_*z*_ *imply that v*_*i*_ = 0 *for i* ∈ 𝒯_*v*_ *and z*_*i*_ = 0 *for i* ∈ 𝒯_*z*_. *It is a CMCS if it contains no smaller cut sets*.

The following lemma relates Definition 1 to the IISs of the second-order cone system, (15).

#### Lemma 1

*Each CMCS 𝒞* = {𝒞_*v*_, 𝒞_*z*_} *for target set* {𝒯_*v*_, 𝒯_*z*_} *under (15b), (15c), (15g), and (15h) corresponds to an IIS of (15)*.

The proof is similar to that of Lemma 1 in [6].

Proof. Consider the CMCS 𝒞 = {𝒞_*v*_, 𝒞_*z*_} for target set {𝒯_*v*_, 𝒯_*z*_}. The definition specifies an infeasible system consisting of (15a)-(15c), (15f)-(15h), and

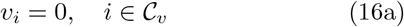

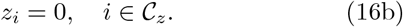

Denote this system Ψ. Ψ is a subsystem of (15) because (16) is a subsystem of *v* = **0** and *z* = **0**. If Ψ is not irreducible, it must contain an IIS. Because the cut set is minimal, the removal of any element from 𝒞_*v*_ or 𝒞_*z*_ makes Ψ feasible. Therefore, any IIS of Ψ must contain (16). Because 𝒞 = {𝒞_*v*_, 𝒞_*z*_} is distinct to the CMCS, any IIS of Ψ is also distinct to the CMCS. D Note the following two implications of Lemma 1: while each IIS corresponds to at most one CMCS, a CMCS can correspond to multiple IISs; and there may be IISs that do not correspond to a CMCS.

We now seek to relate the IISs of (15) to some dual system, which we recall was specified by Farkas’ Lemma in the linear case in [6]. Farkas’ Lemma does not apply to (15) due to the SOC constraint, (15h). There are several extensions to semidefinite systems, e.g., [35], which [9] employs to generalize the results of [7, 8] to semidefinite systems. The dual system in this case is also referred to as the alternative spectrahedron. Because any SOC constraint can be written as a semidefinite constraint, the results of [9] specialize to SOC systems like (15) without modification.

The dual system of (15) is

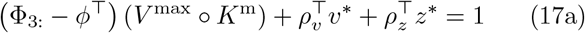

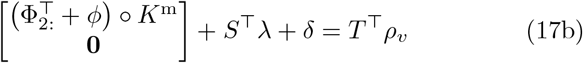

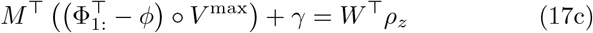

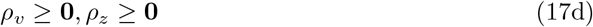

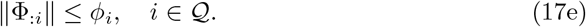

*λ* is the dual variable of (15b); *δ* of *v* = **0**; *γ* of *z* = **0**; *ρ*_*v*_ of (15a); *ρ*_*z*_ of (15f); and (Φ_*i*_, *ϕ*_*i*_), *i* ∈ 𝒬, of the SOC form of (15h).

Theorem 3.2 in [9] states that if (15) is weakly infeasible, then each of its IISs corresponds to an extremal point of (17). More precisely, each IIS determines the nonzero entries of some extremal point of (17). We can thus identify CMCSs by finding the extremal points of (17).

Unfortunately, (17) may have extremal points that do not correspond to an IIS of (15). Theorem 4.1 in [9] provides conditions under which the alternative spectrahedron has a single solution; however, it does not appear to apply in general to (17).

Lemma 3.1 in [9] states that the indices of a minimal cardinality solution of (17) correspond to an IIS of (15). In our case, similar to [6], we seek solutions in which the vectors *δ* and *γ* have minimal cardinality.

In the linear case, there are several different MILPs for solving for IISs. Likewise, one could formulate multiple mixed-integer SOCPs (MISOCP) for finding the IISs of (15); e.g., (9) in [9] can be expressed as an MISOCP when specialized to our problem. The MISOCP below differs from [9] in that we only seek minimal cardinality in *δ* and *γ*, and not the other variables. The portion of the problem corresponding to reaction fluxes is based on the MILP (14) in [36].

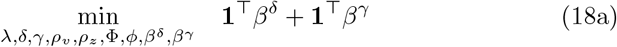

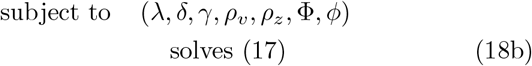

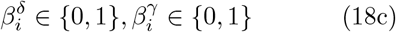

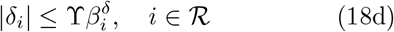

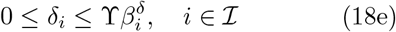

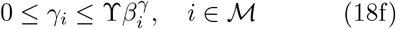

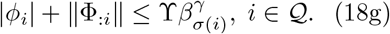

(18c)-(18g) are disjunctive constraints that, for sufficiently large ϒ, either force their left hand sides to be zero or have no effect. The left side of (18g) is the 2-norm of the SOC variable (Φ_:*i*_, *ϕ*_*i*_) [25]. We include this constraint because if 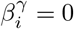, the dual variables associated with both (15h) and (15i) should be zero. Given a solution, the IIS is specified by the entries of *β*^*δ*^and *β*^*γ*^ that are equal to one.

There are a number of further refinements one can make to (18). For example, by weighting the terms in the objective, one can promote cuts with reactions or with metabolites. To exclude either the reactions or metabolites in the target set, or a cut that has already been found, say 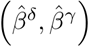, we can add the constraint

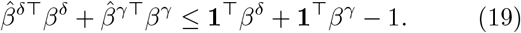

### 4.2. Linear approximation

There are severals reasons why solving (18) might not be the best way to identify CMCSs. First, as discussed, the underlying theoretical results are weaker than in the linear case—an extremal point of the alternative spectrahedron might not correspond to an IIS [9]. Second, MISOCP is less tractable than MILP. And third, given the discrete nature of cut sets, it is not clear that the nonlinearity could not be replaced with something simpler.

The following is a standard linear approximation for when the metabolite concentration is smaller than the Michaelis constant:

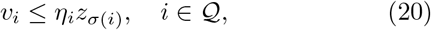

where 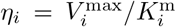. If *z*_*σ*(*i*)_ is in a cut set, i.e., we set it to zero, then *v*_*i*_ ≤ 0 under either (15h) or (20). Definition 1 and Lemma 1 also hold in both cases. Note that while this definition of *η* has precedent, any positive value would serve similarly.

Let (15)* denote (15) with (20) instead of (15h), and let *α* ∈ ℝ^*q*^ be the dual variable of (20). The dual system of (15)* is

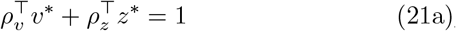

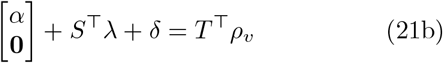

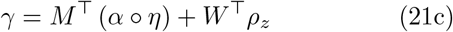

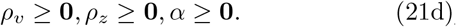

Because (15)* is polyhedral, the results of [7, 8] apply— each IIS of (15)* corresponds to exactly one extreme point of (21). Lemma 1 establishes that each CMCS corresponds to some IIS of (15)*.

Similar to (18), we can identify the IISs by solving the below MILP.

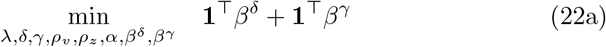

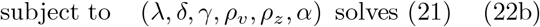

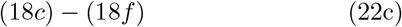

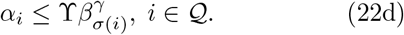

Here (22d) serves the same role as (18g), but is linear because (20) is a linear inequality.

## 5. *Escherichia coli* example

We now apply the tools we have developed to an exam-ple based on the model `e_coli_core` in the BiGG database [37], which corresponds to *Escherichia coli str. K-12 substr. MG1655*. The model has 95 reactions and 72 metabolites.

We take all parameters for the FBA routine (7) from the COBRA Toolbox [12, 13]. We augment the model with Michaelis-Menten kinetics for the reactions listed in Table 1. The values of *K*^m^ and *V*^max^ for each reaction were taken from Table 1 in [38].

**Table 1:**
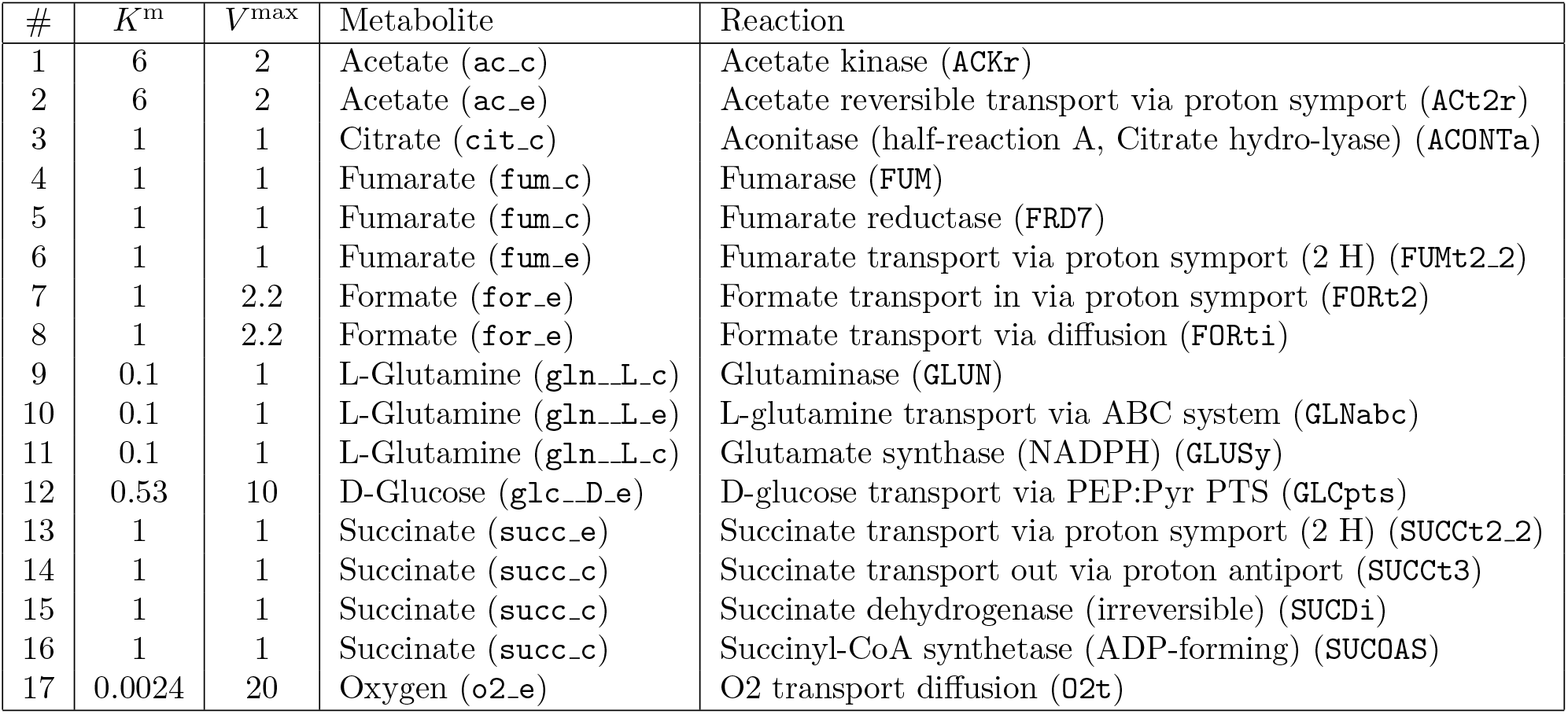
Michaelis-Menten parameters. The units of *K*^m^ and *V*^max^ are mM and mmol/g dw/h, and the values are from [38]. The metabolites and reactions are listed along with their ID in the BiGG database [37].

All optimizations were carried out in Python using CVXPy [39] and the solver Gurobi [40]. All figures were made with Matplotlib [41].

### 5.1. CFBA

We first apply CFBA. We solve the alternative formulation described in Section 3.2.1, which is identical to (9) except in two respects.

- The objective is −**1**^⊤^*z*, which corresponds to minimizing the sum of the concentrations of the metabolites in Table 1.
- We add the constraint *v*_b_ ≥ Ξ. *v*_b_ is the flux of the biomass reaction, which is `BIOMASS_Ecoli_core_w_GAM` in the BiGG database. Here, instead of maximizing the biomass as in FBA, we constrain it to be above Ξ *>* 0.

The SOCP for CFBA took 0.08 seconds to solve. For comparison, the LP for FBA took 0.007 seconds, roughly an order of magnitude less.

Figure 1 shows the sensitivities of the optimal objective to *V*^max^ and *K*^m^ for three of the reaction in Table 1 for Ξ = 0.1. Given primal and dual CFBA solutions, the sensitivities are computed via (11). Each represents the change in total metabolite usage resulting from a small change in a Michaelis-Menten parameter, given that the biomass reaction flux cannot go below Ξ. Of the reactions that are not shown, 1 and 2 have sensitivities on the order of 10^−4^, and the rest 10^−8^.

**Figure 1:**
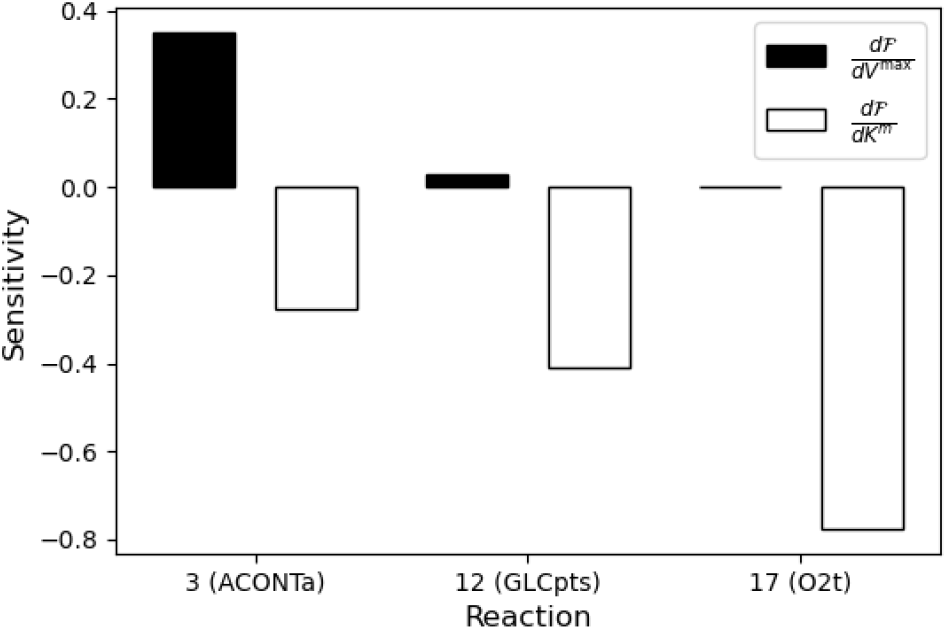
Sensitivities of the optimal CFBA objective, the sum of the metabolite concentrations, to *V*^max^ and *K*^m^ for `ACONTa, GLCpts,` and `O2t`. The lower bound on the biomass reaction flux is Ξ = 0.1.

We can see that the objective is most sensitive to the kinetics of Reaction 3 (`ACONTa`), which depends on the concentration of citrate. This reaction produces H_2_0 for a number of other reactions. Via inspection of the network structure and optimal reaction fluxes, we can see that it also directly enables the reactions aconitase (half-reaction B, Isocitrate hydrolyase) and then isocitrate dehydrogenase (NADP).

The objective is also sensitive to *K*^m^ for Reactions 12 (`GLCpts`) and 17 (`O2t`), which depend on glucose and oxygen, indicating that an increase in *K*^m^ for either reaction will significantly increase the amount of metabolite needed to keep the biomass reaction flux at Ξ.

This indicates that at the optimal solution, Reactions 3, 12, and 17 are successively further to the left on the steeply increasing part of the Michaelis-Menten function; e.g., a slight increase in oxygen would significantly increases the maximum rate of Reaction 17, where as a slightly increase in citrate would moderately increase the maximum rate of Reaction 3. We can conclude that, for this setup, citrate, glucose, and oxygen are the limiting metabolites for biomass production.

Figure 2 shows the concentrations of citrate, glucose, and oxygen as Ξ, the required biomass flux, increases from 0.05 to 0.5. Because these are limiting metabolites, their concentrations increase with Ξ, and do so at a greater than linear rate. The concentrations of citrate and glucose increase rapidly with *ξ* because, at the optimal solution, they are further to the right on flatter part of the Michaelis-Menten function. The concentration of oxygen increases least rapidly because, as described above, a slight increase dramatically increases the reaction rate.

**Figure 2:**
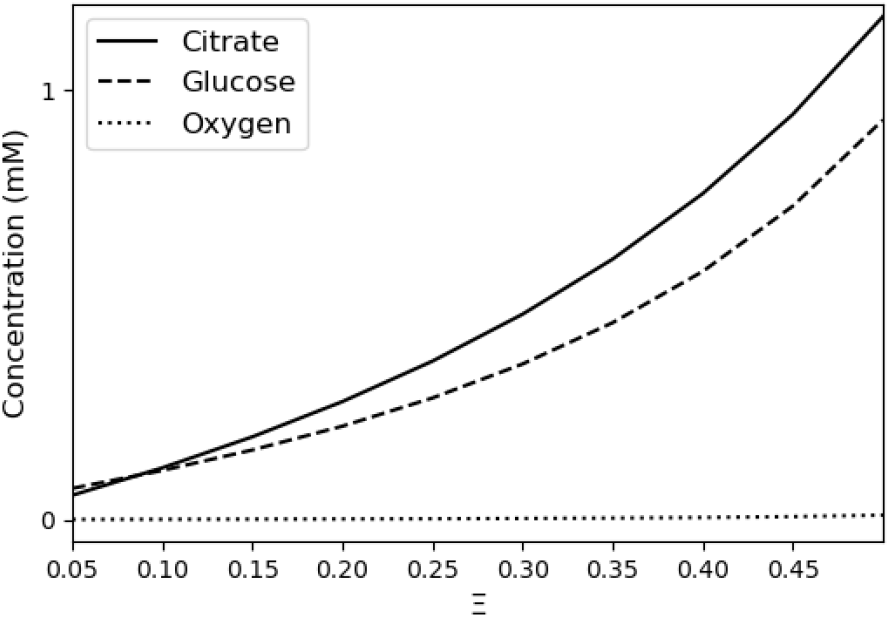
Optimal concentrations of citrate, glucose, and oxygen as a function of the lower bound on the biomass reaction flux, Ξ, in CFBA.

These concentrations are consistent with reported ranges. When Ξ = 0.5, the predicted concentration of citrate is 1.17 mM, within the range of 1.1 to 3.5 mM reported in Supplementary Table 3 of [42]. *Escherichia coli* ‘s metabolism can function over a wide range of oxygen concentrations. When Ξ = 0.5, the predicted oxygen concentration is 0.01 mM, which, for example, falls well within the range depicted in Figure 4A of [43].

### 5.2. Dynamic CFBA

We now test dynamic CFBA by solving (14). Our secondary goal in this section is to understand the scalability of CFBA and its dynamic extension, which we do by varying the time horizon in (14), *τ*.

The objective, (14a), is to maximize the biomass concentration in the last period, *z*_b_(*τ*), as in equation (6b) in Case 2 of [5]. The initial biomass concentration is *z*_b0_ = 0.001, and the remaining elements of *z*_0_ are ones. The biomass concentration evolves as

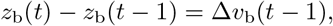

where *v*_b_(*t*), as described in Section 5.1, is the biomass reaction flux.

As described in Section 3.3, we approximate the Michaelis-Menten kinetics with the Contois function, parameterized using the values in Table 1. This means using constraints (14h) and (14i) (and not (14f) or (14g)).

We solved (14) for time horizons ranging from *τ* = 50 to *τ* = 500. In each instance the time step was Δ = 0.001 hours. Figure 3 shows the time taken by the solver as a function of *τ*. When *τ* = 50, there are 8,473 variables, and when *τ* = 500, there are 84,073 variables. The trend is increasing with problem size, though some smaller instances take longer than larger ones, presumably due to specific problem structure and solver behavior. The longest time taken by the solver was roughly six minutes. This confirms that, by virtue of being SOCPs, CFBA and dynamic CFBA are highly tractable problems.

**Figure 3:**
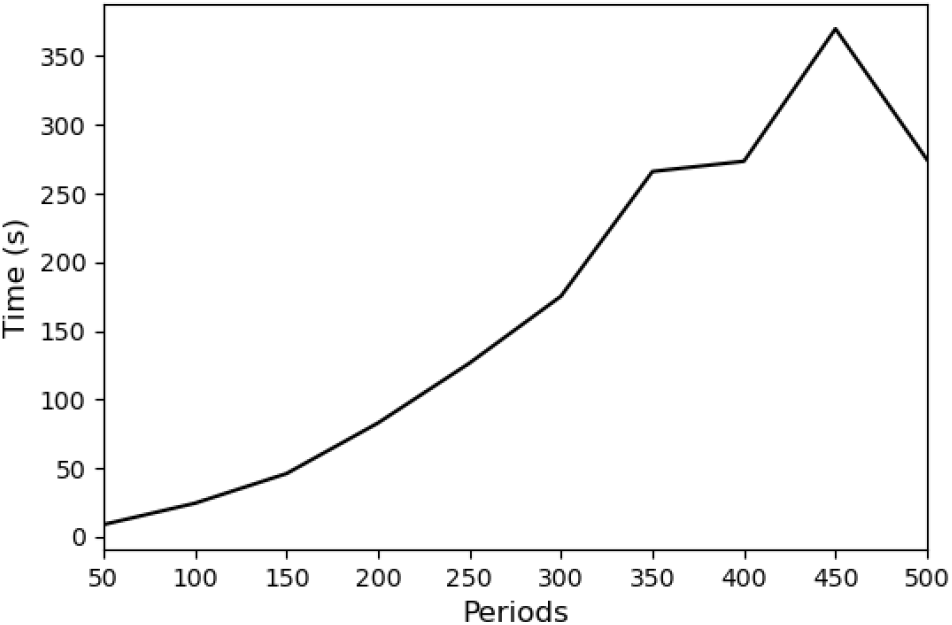
Computation time of dynamic CFBA as a function of the number of periods, *τ*.

**Figure 4:**
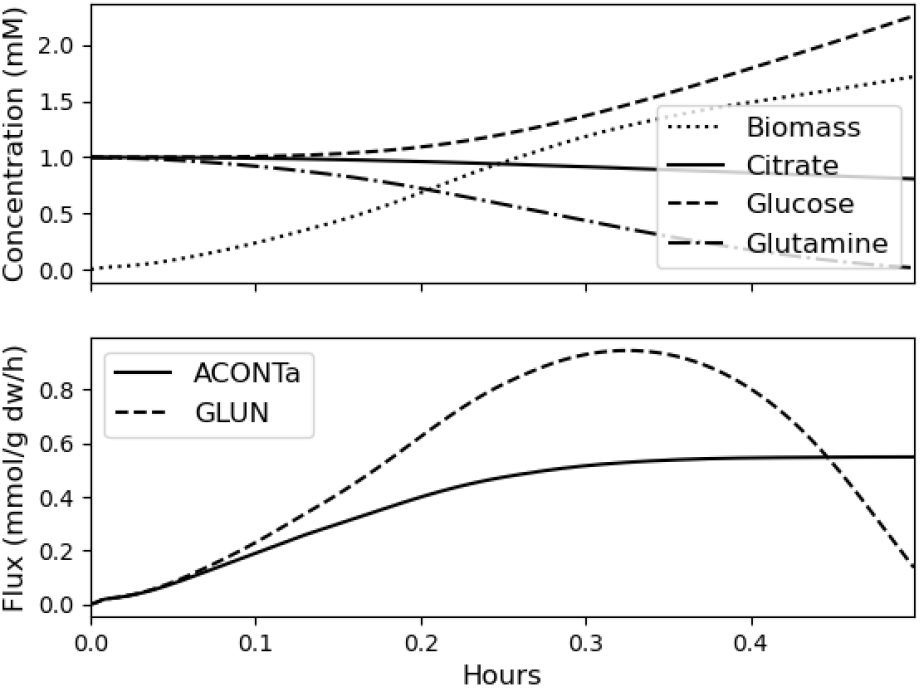
Concentrations (upper) and reactions (lower) through time produced by dynamic CFBA with *τ* = 500 periods.

The upper plot in Figure 4 shows the biomass, citrate, glucose, and glutamine concentrations through time for the case when *τ* = 500. Recall that in Section 5.1, citrate and glucose were two of the metabolites CFBA identified as most important. Glutamine enables the glutaminase reaction, *v*_9_ (`GLUN`), and citrate enables aconitase, *v*_3_ (`ACONTa`), both of which are shown in the lower plot.

The biomass concentration increases from near zero, first exponentially, and then more gradually—this is a standard behavior. The glutaminase reaction first increases as the biomass concentration increases, and then decreases as the concentration of glutamine drops. The aconitase reaction also increases as biomass concentration increases and then levels out.

These observations point to several potential refinements of dynamic CFBA. First, to induce more complicated behaviors like loss or death of biomass, one must include more complicated exogenous conditions; e.g., in [38] there are five different metabolic phases, some of which have distinct exogenous inputs.

A second potential shortcoming is that dynamic CFBA assumes too much foresight, in that the entire trajectory of 500 periods is optimized at once, and with full knowledge of future exogenous inputs. Full foresight is consistent with the dynamic optimization approach, as described in Section 3.3. However, it is the static optimization approach that finds more usage today, wherein optimization only oc-curs within individual time periods. A potential remedy is to implement dynamic CFBA with a shorter horizong in receding horizon fashion [44]. This would both limit foresight and enhance scalability.

### 5.3. CMCS analysis

We compute CMCSs of our example by solving (18), an MISOCP, and (22), an MILP. In addition to the setup at the start of this section, we must also specify target sets, 𝒯_*v*_ and 𝒯_*z*_, in the form of (15a) and (15f).

Table 2 lists reaction ‘knockouts’ and the corresponding CMCSs found by (18) and (22), specified by their IDs in the BiGG database [37]. The target set in each case consists of constraining the flux to be greater than one. Note that we only list the first CMCSs found by the solver, that one target set can have many CMCSs, and that a given CMCS can be a solution for both (18) and (22).

**Table 2:**
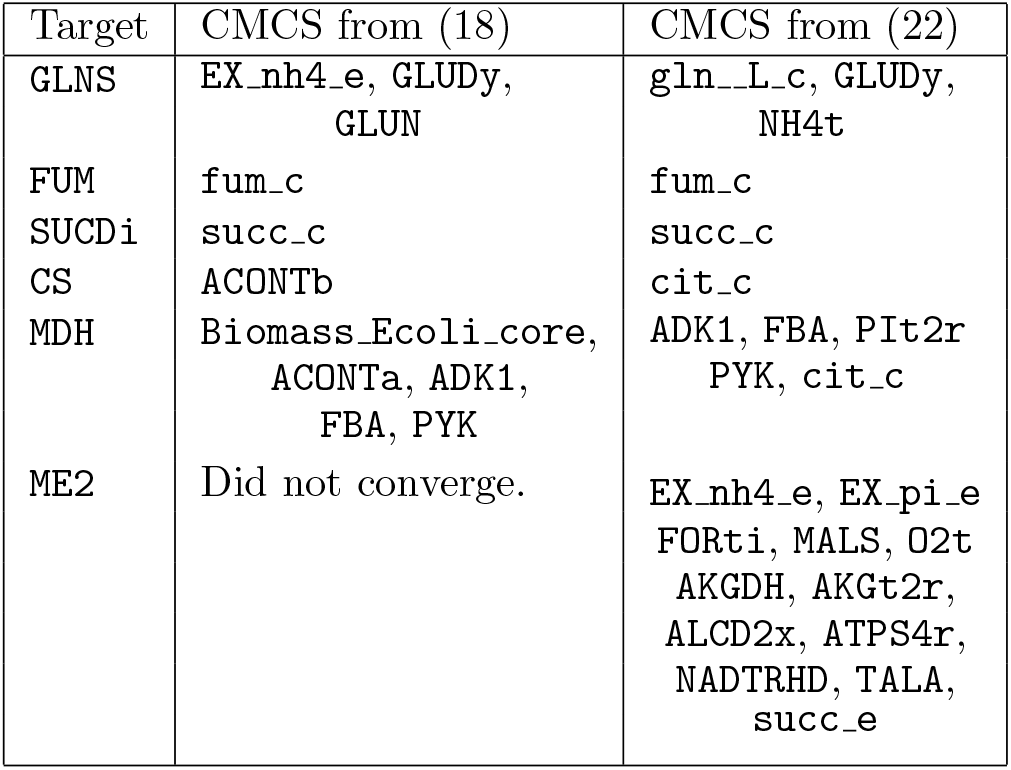
Target knockout reactions and the CMCSs found via (18) and (22).

Whereas an MCS contains only critical reactions, the CMCSs in Table 2 consist of reactions, metabolites, or combinations of both. The prevalence of metabolites in the CMCSs depends heavily on which reactions are limited by Michaelis-Menten kinetics. For instance, `fum_c` and `succ_c` are both CMCSs because they limit `FUM` and `SUCDi`, among other reactions.

The computation time in all but the final case was under one second. In the last case, (22) took six seconds to solve, and (18) was still not solved after an hour. This highlights the fact that MILP and MISOCP are NP-hard, and similar instances of a problem can take very different times to solve.

Problem (22) often has the same or similar solutions to (18) and, due to the higher tractability and maturity of MILP, is substantially easier to solve in some cases. For these reasons, (22) appears to be more practical than (18).

## 6. Conclusion

By representing the Michaelis-Menten kinetics as a second-order cone constraint, we can add metabolite concentrations to standard models of metabolic networks without losing much tractability. This has enabled us to formulate several new tools: conic flux balance analysis, dynamic conic flux balance analysis, and conic minimal cut set analysis. In our numerical examples, we demonstrated that each of these new problems is tractable and can provide insight into both reaction fluxes and metabolite concentrations.

There are several directions for future work. We believe that there are numerous potential applications to the many different organisms there are. Such studies could both provide new insights into metabolic networks and further clarify when these tools are appropriate. A starting point for this is applying them to larger metabolic network models. This entails augmenting more existing models with Michaelis-Menten parameter data. There is also room for methodological advancements. For example, a receding horizon implementation [44] could make dynamic conic flux balance analysis both more realistic and tractable. The SOC representation of Michaelis-Menten could be incorporated into flux variability analysis [45] so as to find near optimal ranges of both fluxes and metabolites. There is certainly more to understand about the basic geometry of our setup, which, though convex, is not amenable to many of the techniques used to analyze polyhedral models.

## Code availability

The code for the examples is available at: https://github.com/JAT38/conic-metabolic.

## Acknowledgements

We thank Professor Christopher Lawson for helpful discussion.

## Appendix A Second-order cone programming

We here provide cursory background on SOCP, and refer the reader to [25] and [4] for in-depth coverage. Let *A* ∈ ℝ^*m*×*n*^, *b* ∈ ℝ^*m*^, *c* ∈ ℝ^*n*^, and *d* ∈ ℝ. A standard form SOC constraint on the variable *x* ∈ ℝ^*n*^ is written

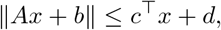

where the left-hand side is the two-norm. If *A* = **0**, this reduces to a linear constraint.

A commonly occurring constraint with SOC form is the hyperbolic constraint 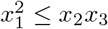. It can be written

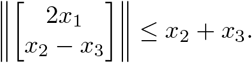

We must also require *x*_2_ ≥ 0 and *x*_3_ ≥ 0.

An optimization problem with a linear objective and *p* SOC constraints is an SOCP, and can in general be written

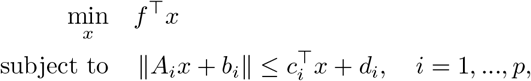

where, 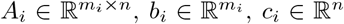, and *d*_*i*_ ∈ ℝ. Note that the number of rows in *A*_*i*_ and *b*_*i*_ can be different for each *i*. SOCP is a generalization of LP because any linear constraint can be written as an SOC constraint.

The SOC is self-dual, i.e., its dual cone is the SOC. The dual of an SOCP is therefore also an SOCP. The dual of the above SOCP is

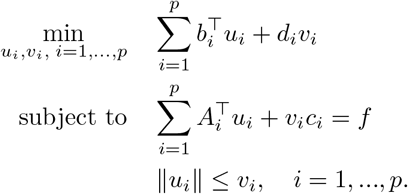

Strong duality is attained if a constraint qualification holds, e.g., the existence of a Slater point. In this case the two optimizations have the same objective.

